## Supplementary material for "The larva and adult of *Helicoverpa armigera* use differential gustatory receptors to sense sugars": Figure supplements for Figure 1-7

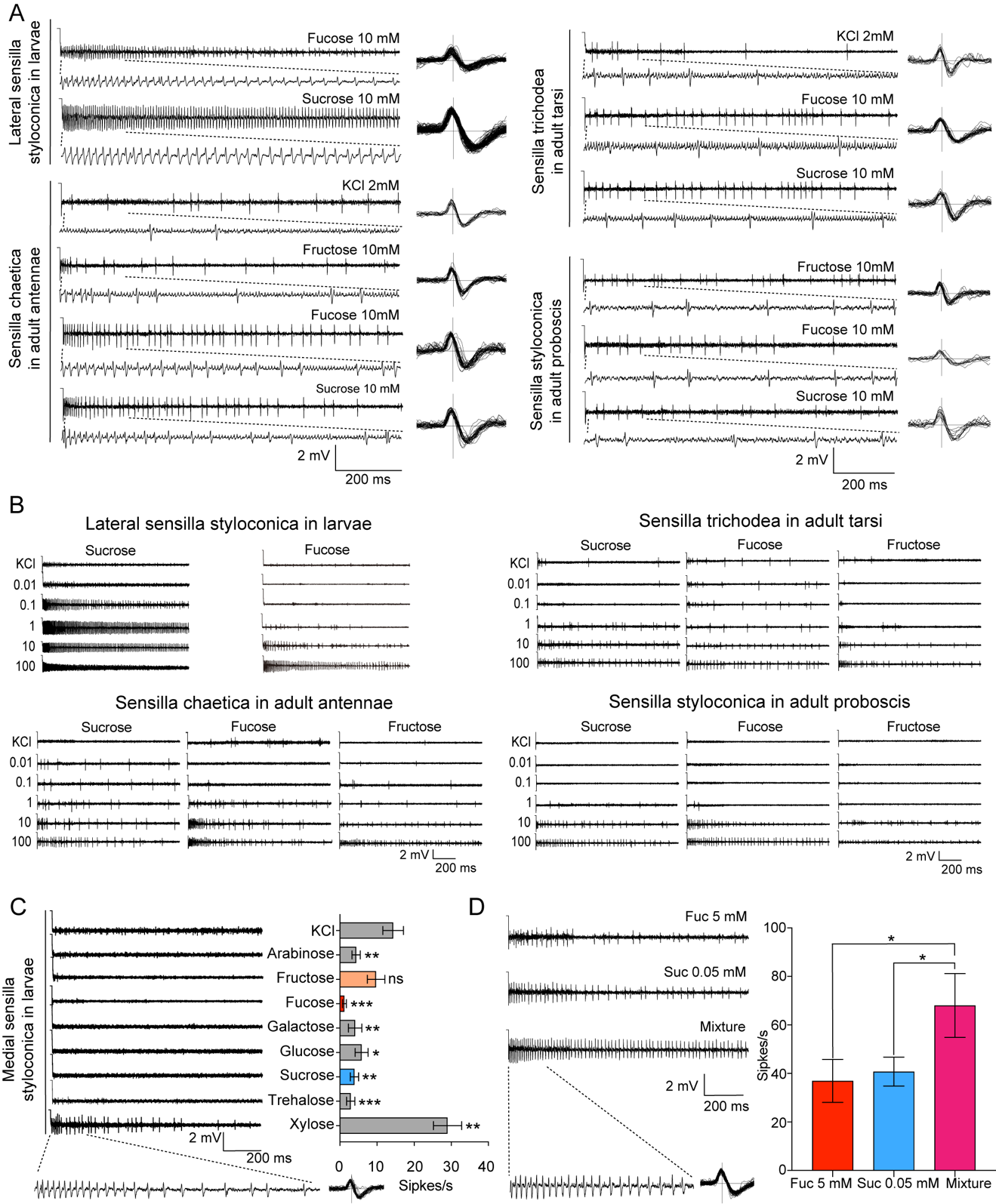

**Figure 1—figure supplement 1. Electrophysiological responses of contact chemosensilla in *Helicoverpa armigera* to sugar compounds.** **(A)** The representative spike traces and waveforms of lateral sensilla styloconica in larval maxillary galea, sensilla chaetica in adult antennae, sensilla trichodea in adult tarsi, and sensilla styloconica in adult proboscis response to test compounds. **(B)** Representative dose–response spike traces of lateral sensilla styloconica in larval maxillary galea, sensilla chaetica in female adult antennae, sensilla trichodea in female adult tarsi and sensilla styloconica in female adult proboscis to sucrose, fucose, and fructose. **(C)** The representative spike traces of the responses of medial sensilla styloconica in larval maxillary galea to eight sugars at 10 mM (left), and quantifications of the firing rates (right) (*n =* 13). **(D)** The representative spike traces of the responses of lateral sensilla styloconica in larval maxillary galea (left), and quantifications of the firing rates to 0.05 mM sucrose, 5 mM fucose, and the mixture of those (right) (*n =* 6). **(A–B)** The dotted lines indicate the spike trace of the first 200 ms. **(C–D)** Data are mean ± SEM; * *P* < 0.05; ** *P* < 0.01; *** *P* < 0.001; ns indicates no significance (independent-samples *t* test).

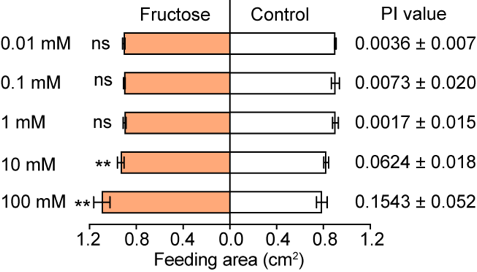

**Figure 2—figure supplement 1. Feeding responses and the PI value of 5^th^ instar larvae of *Helicoverpa armigera* to different concentrations of fructose painted on the cabbage leaf discs in two-choice tests.** Data are mean ± SEM; ** *P* < 0.01; ns indicates no significance (paired *t* test, *n =* 20).

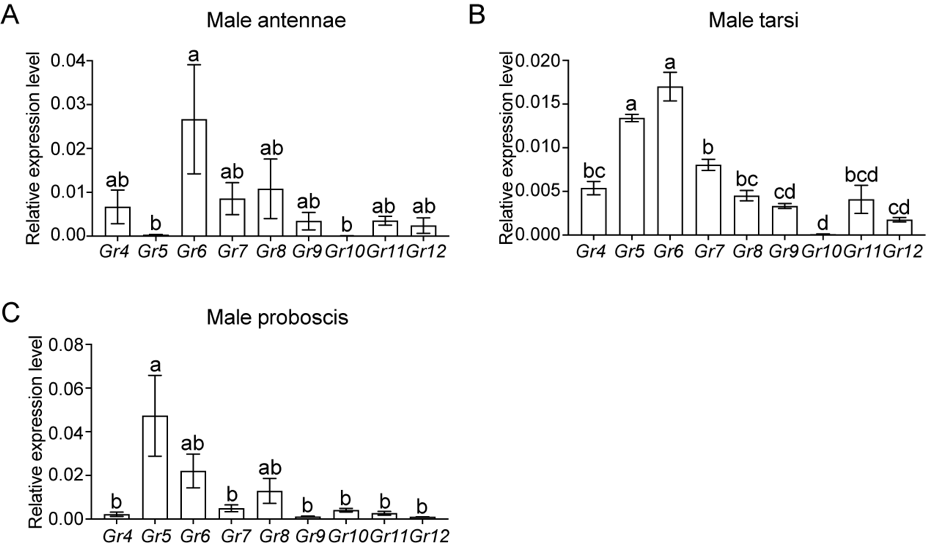

**Figure 3—figure supplement 1. Expression patterns of sugar GRs in taste organs of *Helicoverpa armigera* male adults. (A)** Relative expression levels of sugar GRs in male antennae determined by qRT-PCR. **(B)** Relative expression levels of sugar GRs in male tarsi. **(C)** Relative expression levels of sugar GRs in male proboscis. **(A–C)** Data are mean ± SEM. One-way ANOVA was used, and different letters labeled indicate significant difference (Tukey’s HSD test, *P* < 0.05, *n =* 3).

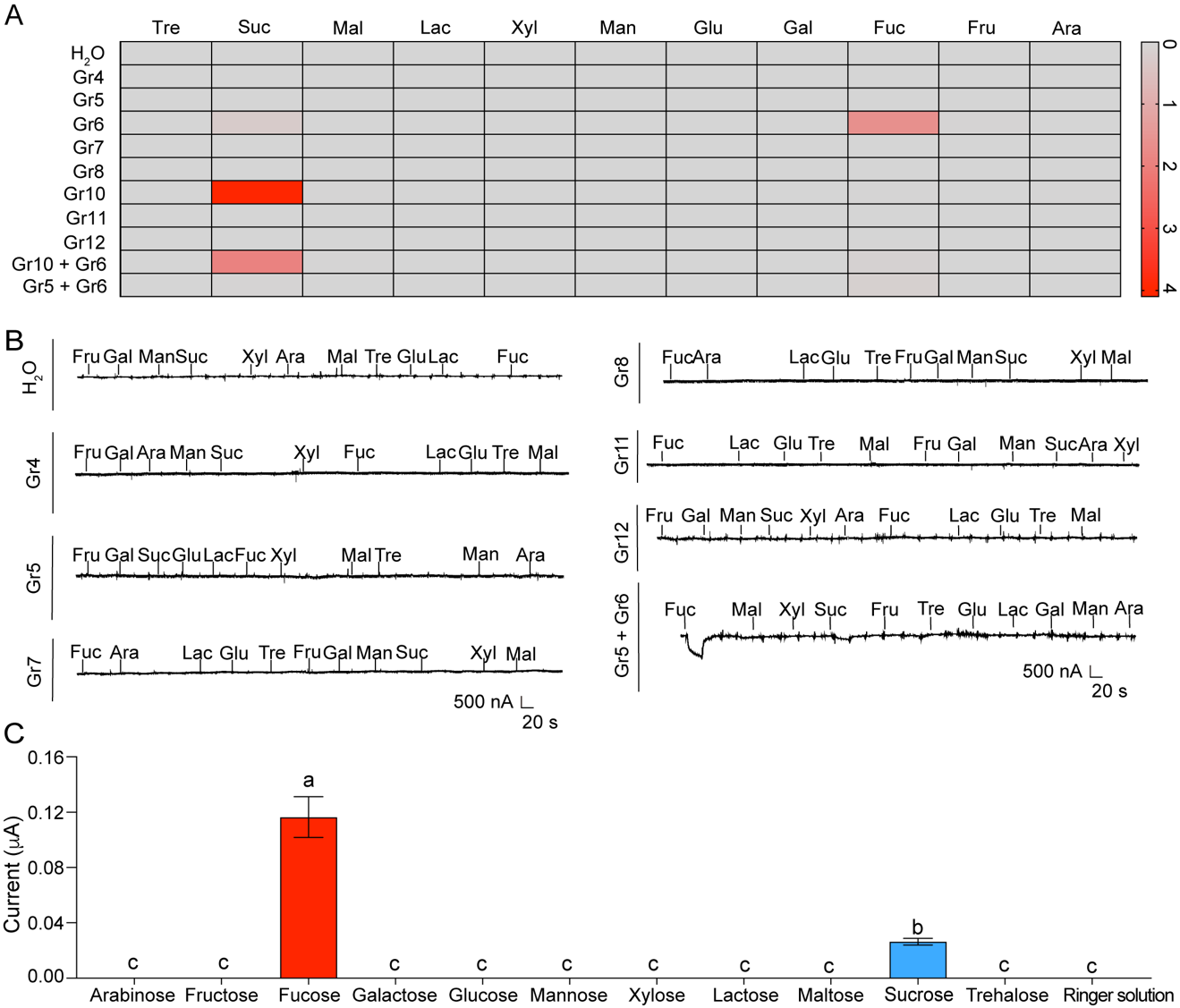

**Figure 4—figure supplement 1. The inward current responses and representative traces of *Xenopus* oocytes expressing sugar GRs of *Helicoverpa armigera*.** **(A)** Heat-map signal indicates the mean of the responses to eleven sugars at 100 mM of the oocytes expressing *H. armigera* sugar GRs (H_2_O: mannose, *n =* 5; other sugar compounds: *n =* 6. Gr4: arabinose, *n =* 3; other sugar compounds: *n =* 7. Gr5: *n =* 7. Gr6: mannose, *n =* 15; other sugar compounds: *n =* 20. Gr7: *n =* 5. Gr8: *n =* 5. Gr10: maltose, *n =* 13; other sugar compounds: *n =* 14. Gr11: *n =* 6. Gr12: *n =* 6. Gr10 + Gr6: *n =* 6. Gr5 + Gr6: *n =* 10). Gray and red signals represent lower and higher response levels, respectively. **(B)** The representative two-electrode voltage-clamp traces of oocytes expressing Gr4, Gr5, Gr7, Gr8, Gr11, and Gr5 + Gr6 to eleven sugars at 100 mM and the control (H_2_O). **(A-B)** Ara: arabinose; Fru: fructose; Fuc: fucose; Gal: galactose; Glu: glucose; Lac: lactose; Mal: maltose; Man: mannose; Suc: sucrose; Tre: trehalose; Xyl: xylose; Rin: ringer solution. **(C)** The responses of the oocytes expressing Gr5 + Gr6 to sugars at 100 mM (*n* = 10). Data (mean ± SEM) were analyzed by one-way ANOVA with Tukey’s HSD test, and different letters labeled indicate significant differences (*P* < 0.05).

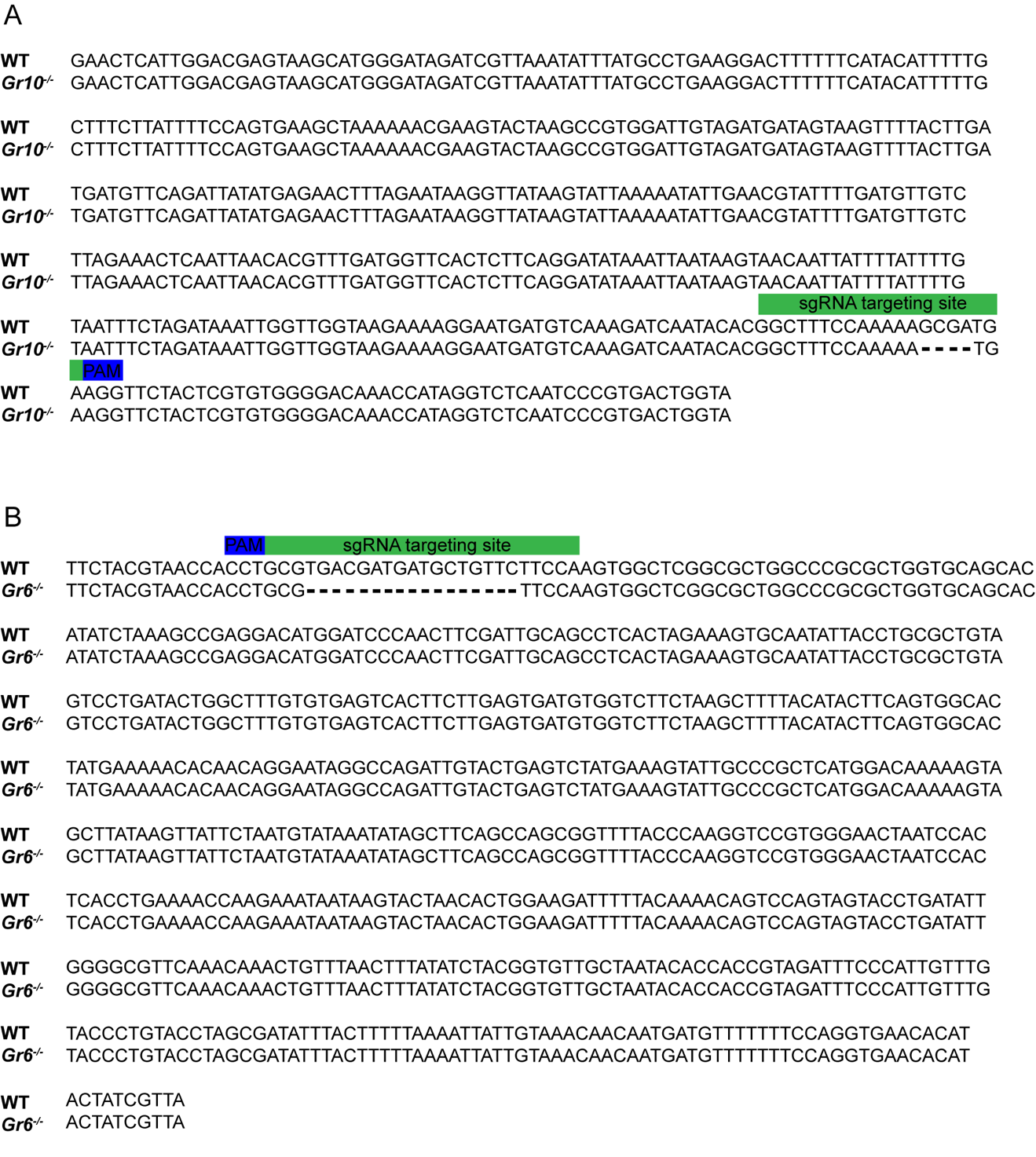

**Figure 5—figure supplement 1. Confirmation of the deletion of *Gr10* and *Gr6* in *Gr10^-/^*^-^ and *Gr6^-/-^* mutants at mRNA level**. **(A)** The alignment of the nucleic acid sequences based on *Gr10* transcripts in WT and *Gr10^-/-^*. **(B)** The alignment of the nucleic acid sequences based on *Gr6* transcripts in WT and *Gr6^-/-^*. The green rectangle corresponds to the position of the sgRNA targeting site, the blue rectangle corresponds to the position of the PAM site.

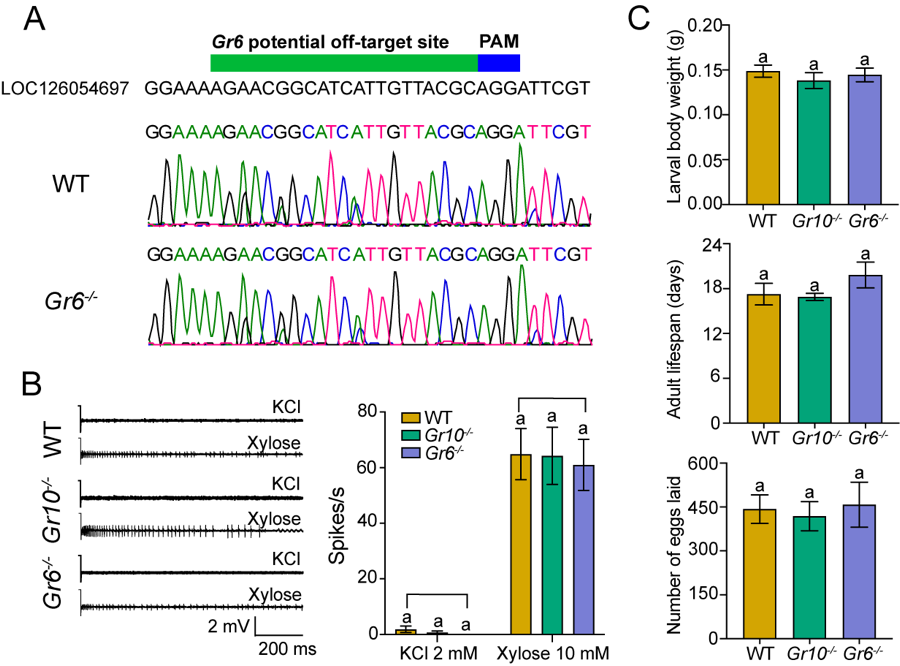

**Figure 5—figure supplement 2. The potential off-target effects detection. (A)** Representative chromatograms of potential off-target PCR products obtained from the wild type (WT) and *Gr6^-/-^*. **(B)** The representative spike traces (the left) and quantifications of the firing rate (the right) of medial sensilla styloconica in larval maxillary galea to 2 mM KCl and 10 mM xylose (*n =* 8). **(C)** The larval body weight at the beginning of the fifth instar (the upper, *n* = 16), the adult lifespan (the middle, *n* = 16), and the number of eggs laid (the lower, *n* = 5). **(B–C)** Data are mean ± SEM. Data were analyzed by one-way ANOVA with Turkey’s HSD test, and different letters labeled indicate significant differences (*P* < 0.05).

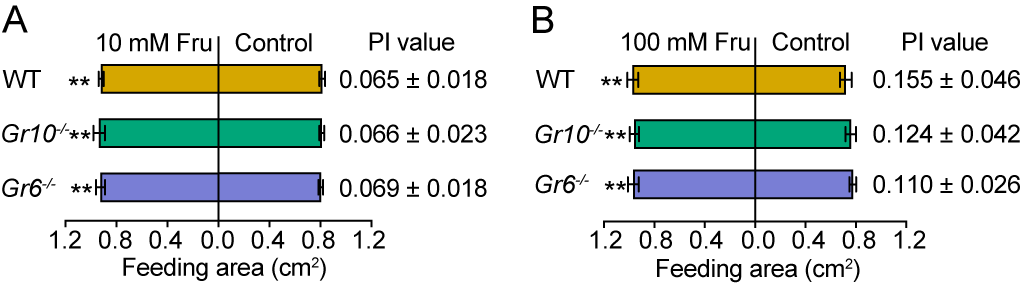

**Figure 7—figure supplement 1. Feeding responses and the PI value of 5^th^ instar larvae of WT, *Gr10^-/-^* and *Gr6^-/-^* of *Helicoverpa armigera* to fructose painted on the cabbage leaf discs in two-choice tests. (A)** Feeding area and the PI value to 10 mM fructose. **(B)** Feeding area and the PI value to 100 mM fructose. Data are mean ± SEM, *n =* 20, paired *t* test, ** *P* < 0.01. Fru, fructose.
