## Appendix for "The larva and adult of *Helicoverpa armigera* use differential gustatory receptors to sense sugars"

**Template:**

| **Key Resources Table** | | | | |
| --- | --- | --- | --- | --- |
| **Reagent type (species) or resource** | **Designation** | **Source or reference** | **Identifiers** | **Additional information** |
| gene (*Helicoverpa armigera*) | Gr4 | GenBank | OP251144 | More details about this gene see Table S1. |
| gene (*H. armigera*) | Gr5 | GenBank | OP251145 | More details about this gene see Table S1 |
| gene (*H. armigera*) | Gr6 | GenBank | OP251146 | More details about this gene see Table S1 |
| gene (*H. armigera*) | Gr7 | GenBank | OP251147 | More details about this gene see Table S1 |
| gene (*H. armigera*) | Gr8 | GenBank | OP251148 | More details about this gene see Table S1 |
| gene (*H. armigera*) | Gr9 | GenBank | XM_049843199 | More details about this gene see Table S1 |
| gene (*H. armigera*) | Gr10 | GenBank | OP251149 | More details about this gene see Table S1 |
| gene (*H. armigera*) | Gr11 | GenBank | OP251150 | More details about this gene see Table S1 |
| gene (*H. armigera*) | Gr12 | GenBank | OP251151 | More details about this gene see Table S1 |
| strain, strain background (include species and sex here) | N/A | N/A | N/A | N/A |
| genetic reagent (include species here) | N/A | N/A | N/A | N/A |
| cell line (include species here) | N/A | N/A | N/A | N/A |
| transfected construct (include species here) | N/A | N/A | N/A | N/A |
| biological sample (include species here) | N/A | N/A | N/A | N/A |
| antibody | N/A | N/A | N/A | N/A |
| recombinant DNA reagent | N/A | N/A | N/A | N/A |
| sequence-based reagent | N/A | N/A | N/A | N/A |
| peptide, recombinant protein | N/A | N/A | N/A | N/A |
| commercial assay or kit | RNeasy Plus Universal Mini Kit | Qiagen | Cat# 73404 |  |
| commercial assay or kit | Q5 High-Fidelity DNA Polymerase | NEB | Cat# M0491 |  |
| commercial assay or kit | TransStart FastPfu DNA Polymerase | TransGen Biotech | Cat# AP221-01 |  |
| commercial assay or kit | M-MLV reverse transcriptase | Promega | Cat# M1701 |  |
| commercial assay or kit | SYBR Premix Ex Taq | Takara | Cat# RR820 |  |
| commercial assay or kit | mMESSAGE mMACHINE SP6 | Ambion | Cat# AM1340 |  |
| commercial assay or kit | GeneArt^TM^ gRNA Clean Up Kit | Invitrogen | Cat#A29377 |  |
| commercial assay or kit | GeneArt^TM^ gRNA Prep Kit | Invitrogen | Cat#A29377 |  |
| commercial assay or kit | TrueCut^TM^ Cas9 protein 2 | Invitrogen | Cat#A36498 |  |
| commercial assay or kit | Animal Tissue PCR Kit | TransGen Biotech | Cat#AD201-01 |  |
| chemical compound, drug | L - (+) - Arabinose | Sigma-Aldrich | CAS: 5328-37-0 |  |
| chemical compound, drug | D - (-) - Fructose | Sigma-Aldrich | CAS: 57-48-7 |  |
| chemical compound, drug | L - (-) - Fucose | Sigma-Aldrich | CAS: 2438-80-4 |  |
| chemical compound, drug | D - (+) - Galactose | Sigma-Aldrich | CAS: 59-23-4 |  |
| chemical compound, drug | D - (+) - Glucose | Sigma-Aldrich | CAS: 50-99-7 |  |
| chemical compound, drug | D - (+) - Mannose | Sigma-Aldrich | CAS: 3458-28-4 |  |
| chemical compound, drug | D - (+) - Xylose | Sigma-Aldrich | CAS: 58-86-6 |  |
| chemical compound, drug | D - Lactose monohydrate | Sigma-Aldrich | CAS: 64044-51-1 |  |
| chemical compound, drug | D - (+)-Maltose monohydrate | Sigma-Aldrich | CAS: 6363-53-7 |  |
| chemical compound, drug | Sucrose | Sigma-Aldrich | CAS: 57-50-1 |  |
| chemical compound, drug | D - (+) - Trehalose dihydrate | Sigma-Aldrich | CAS: 6138-23-4 |  |
| chemical compound, drug | Sodium chloride | Sigma-Aldrich | CAS: 7647-14-5 |  |
| chemical compound, drug | Potassium chloride | Sigma-Aldrich | CAS: 7447-40-7 |  |
| chemical compound, drug | Magnesium chloride hexahydrate | Sigma-Aldrich | CAS: 7791-18-6 |  |
| chemical compound, drug | HEPES | Sigma-Aldrich | CAS: 7365-45-9 |  |
| software, algorithm | SAPID Tools software version 3.5 | Smith et al., 1990. https://doi.org/10.1093/chemse/15.3.253. |  |  |
| software, algorithm | Autospike 3.7 | Syntech |  |  |
| software, algorithm | MAFFT version 7.455 | Rozewicki et al., 2019. https://doi.org/10.1093/nar/gkz342. |  |  |
| software, algorithm | trimAI version 1.4 | Capella-Gutierrez et al., 2009. https://doi.org/10.1093/bioinformatics/btp348. |  |  |
| software, algorithm | IQ-tree version 6.8 | Nguyen et al., 2015. https://doi.org/10.1093/molbev/msu300. | iqtree.org |  |
| software, algorithm | pCLAMP software version 10.4.2.0 | Axon Instruments Inc. | RRID:SCR_011323 |  |
| software, algorithm | SnapGene^®^ software version 4.3.8 | Insightful Science | snapgene.com |  |
| software, algorithm | SeqMan software version 7.1 | DNASTAR | dnastar.com |  |
| software, algorithm | SPSS 20 | IBM | ibm.com |  |
| software, algorithm | GraphPad Prism 8.2.1 | Dotmatics | graphpad.com |  |
| other | Primers for full-length cloning of GRs | This paper |  | see Appendix—Table 2 |
| other | Primers for qRT-PCR | This paper |  | see Appendix—Table 3 |
| other | Primers for *Xenopus* oocytes expression system | This paper |  | see Appendix—Table 4 |
| other | Primers for experiments of mutant strains establishment | This paper |  | see Appendix—Table 5 |

**Appendix—Table 1. GenBank accession numbers for sugar gustatory receptors used in this study**

| **Gene** | **Sequence** | **Xu et al., 2017** | **Pearce et al., 2017** | **In this study** |
| --- | --- | --- | --- | --- |
| *Gr4* | Nucleotide | KY806282.1 | XM_021327184.1 | OP251144 |
|  | Protein | ASW18693.1 | XP_021182859.1 |  |
| *Gr5* | Nucleotide | KY806283.1 | XM_021345811.1 | OP251145 |
|  | Protein | ASW18694.1 | XP_021201486.1 |  |
| *Gr6* | Nucleotide | KY806284.1 | XM_021327785.1 | OP251146 |
|  | Protein | ASW18695.1 | XP_021183460.1 |  |
| *Gr7* | Nucleotide | KY806285.1 | XM_021325184.1 | OP251147 |
|  | Protein | ASW18696.1 | XP_021180859.1 |  |
| *Gr8* | Nucleotide | KY806286.1 | XM_021326422.1 | OP251148 |
|  | Protein | ASW18697.1 | XP_021182097.1 |  |
| *Gr9* | Nucleotide |  | XM_049843199 |  |
|  | Protein |  | XP_049699156 |  |
| *Gr10* | Nucleotide | KY806287.1 | XM_021325160.1 | OP251149 |
|  | Protein | ASW18698.1 | XP_021180835.1 |  |
| *Gr11* | Nucleotide | KY806288.1 | XM_021346090.1 | OP251150 |
|  | Protein | ASW18699.1 | XP_021201765.1 |  |
| *Gr12* | Nucleotide | KY806289.1 | XM_021325173.1 | OP251151 |
|  | Protein | ASW18700.1 | XP_021180848.1 |  |

**Appendix—Table 2.** **The primer sequences used in PCR for full-length cloning of GRs**

| **Primer name** | **Primer sequences (5’–3’)** |
| --- | --- |
| *Gr4*F | ATGGAAATTAAGCTGTGTAAACTATTTG |
| *Gr4*R | CTACTTATCATATTGTATTAAAACTAAT |
| *Gr5*F | ATGCAAAATGGATGGAACAACGTTATC |
| *Gr5*R | CTACTTAATAACACCACTGACAGACTC |
| *Gr6*F | ATGGGGCAAACAAGTTTTAGAAG |
| *Gr6*R | TTAAGTAGCTAGTGTTGTAATAGTG |
| *Gr7*F | ATGTCAAGTCGAGGGTTCGGTCAAT |
| *Gr7*R | CTATACTCCAATAGCATTCTTAGTC |
| *Gr8*F | ATGTCGAGTAAGGAATTCAAACAATTTTTG |
| *Gr8*R | TTAAGATGTTAGTAGCATTTAATGCTATCC |
| *Gr9*F | ATGTTGTGGATAGAAACTC |
| *Gr9*R | TCAGTTATCGTATCTTTGGAATTG |
| *Gr10*F | ATGGAATATGGGTTAGATGCTAAAATTTC |
| *Gr10*R | CTAGTTCTTGTTCAATTCGTCGCTGCTGA |
| *Gr11*F | ATGCCATCGAAACTATTTCTCAAGACAT |
| *Gr11*R | TTATCTGTTGAACTGTAGCAACACCAGT |
| *Gr12*F | ATGAAAGTCCACCAACTGACGATGAGAA |
| *Gr12*R | TTAACTGTAACTGTGCGAAGAAATATTT |

F: forward strand; R: reverse strand.

**Appendix—Table 3. The primer sequences used for qRT-PCR**

| **Primer name** | **Primer sequences (5’–3’)** |
| --- | --- |
| *Gr4*F | GCGCATGGTGGATCGAAATC |
| *Gr4*R | CAACACTACACTGCAGGCGCG |
| *Gr5*F | AGATGCAAGCCAGAAATTGGA |
| *Gr5*R | CTTGGATGCAGTGAGGGAGA |
| *Gr6*F | TTCTGGAGAGCTACCCGTGA |
| *Gr6*R | GGAGTGCATTCTCCTGTGCC |
| *Gr7*F | GCAAAGTGGACGATGCCATC |
| *Gr7*R | GTAGGCTGTCTTGATAGGCTC |
| *Gr8*F | TACGTGAAGCAAGCGATGCTG |
| *Gr8*R | ATCTGTGGACATCCCTTGCG |
| *Gr9*F | AGAGGGCAAAAGAACCGAGG |
| *Gr9*R | AGCTGCCGCGAGAATATCTC |
| *Gr10*F | AAGGGAGGATTACACACGCG |
| *Gr10*R | TCAGGTAAGCGGGGTAGGAAT |
| *Gr11*F | GCGGCCTCCATAGTACTGTC |
| *Gr11*R | TCCGGCTAGTAGGACAAGTGA |
| *Gr12*F | CGAAACAATGGCCGAGGTT |
| *Gr12*R | GTGCCATAGCAACGTAAGCG |
| *18S*-F | CGTTGCTGGGAAGTTGACCA |
| *18S*-R | CTTCCGCAGGTTCCCCTACG |

F: forward strand; R: reverse strand.

**Appendix—Table 4. The primer sequences used in cDNA synthesis for *Xenopus* oocytes expression system**

| **Primer name** | **Primer sequences (5’–3’)** |
| --- | --- |
| Gr4 (BamHI) F | *CG*GGATCC**GCCACC**ATGGAAATTAAGCTGTGTAAACTATTTG |
| Gr4 (XhoI) R | *CC*CTCGAGCTACTTATCATATTGTATTAAAACTAAT |
| Gr5 (BamHI) F | GGATCC**GCCACC**ATGCAAAATGGATGGAACAACGTTATC |
| Gr5 (XhoI) R | CTCGAGCTACTTAATAACACCACTGACAGACTC |
| Gr6 (EcoRI) F | GAATTC**GCCACC**ATGGGGCAAACAAGTTTTAGAAGAAATATG |
| Gr6 (Xbal) R | TCTAGATTAAGTAGCTAGTGTTGTAATAGTGTG |
| Gr7 (EcoRI) F | *CG*GAATTC**GCCACC**ATGTCAAGTCGAGGGTTCGGTCAAT |
| Gr7 (XbaI) R | *GC*TCTAGACTATACTCCAATAGCATTCTTAGTC |
| Gr8 (StuI) F | *GA*AGGCCT**GCCACC**ATGTCGAGTAAGGAATTCAAACAATTTTTG |
| Gr8 (XbaI) R | *GC*TCTAGATTAAGATGTTAGTAGCATTTAATGCTATCC |
| Gr10 (BamHI) F | GGATCC**GCCACC**ATGGAATATGGGTTAGATGCTAAAATTTC |
| Gr10 (XhoI) R | CTCGAGCTAGTTCTTGTTCAATTCGTCGCTGCTGA |
| Gr11 (BamHI) F | *CG*GGATCC**GCCACC**ATGCCATCGAAACTATTTCTCAAGACAT |
| Gr11 (XbaI) R | *GC*TCTAGATTATCTGTTGAACTGTAGCAACACCAGT |
| Gr12 (BamHI) F | *CG*GGATCC**GCCACC**ATGAAAGTCCACCAACTGACGATGAGAA |
| Gr12 (XbaI) R | *GC*TCTAGATTAACTGTAACTGTGCGAAGAAATATTT |

The italic sequences are protective bases, underline sequences are restriction enzymes, bold sequences are Kozak sequences. F: forward strand; R: reverse strand.

**Appendix—Table 5. The primer sequences used in experiments of mutant strains establishment**

| **Primer name** | **Primer sequences (5’–3’)** |
| --- | --- |
| **For sgRNA synthesis** |  |
| sgGr6F | TAATACGACTCACTATAGGAACAGCATCATCGTCACGC |
| sgGr6R | TTCTAGCTCTAAAACGCGTGACGATGATGCTGTTC |
| sgGr10F | TAATACGACTCACTATAGGGCTTTCCAAAAAGCGATGA |
| sgGr10R | TTCTAGCTCTAAAACTCATCGCTTTTTGGAAAGCC |
| **For mutation detection** |  |
| mGr6F | CACCAGTAATATTCTACGTAACCACC |
| mGr6R | AAACCTTCGTAAGTATCCATTCCTGA |
| mGr10F | CGAAGGAATTAGAAAGTATCAATTTC |
| mGr10R | CTTTCTGTAGTATACCAGTC |
| **For deletion confirmation** |  |
| dGr6F | TACTCGTTGCTGTCGCTCAT |
| dGr6R | CCCGCAAATGCTGATAATAGGG |
| dGr10F | GGGTTAGATGCTAAAATTTCGAAGGA |
| dGr10R | CATCAACGACGGCCATTTTGA |
| **For off-target detection** |  |
| Gr6F | ATAGAACGTGAACGGAGGGC |
| Gr6R | CGTCAGACCCGTAGGTATGC |

F: forward strand; R: reverse strand.

**Appendix—Table 6. Summary of the CRSPR/Cas9 directed mutation rates from G0 to G2**

|  | **Injected embryo** | **Chimeras rate in G0** | ***Gr^n/-^* rate in G1** | ***Gr^-/^*^-^ rate in G2** |
| --- | --- | --- | --- | --- |
| ***Gr10^-/-^*** | 732 | 15/125(12%) | 24/57(42.1%) | 53/254(20.8%) |
| ***Gr6^-/-^*** | 695 | 12/132(9.1%) | 19/66(28.7%) | 42/200(21%) |
